# DOWN state in the anterior cingulate and prelimbic areas in rats during immobility

**DOI:** 10.1101/847913

**Authors:** Anatol Bragin, Lin Li, Sotiris C. Masmanidis, Jerome Engel

## Abstract

**Background:** UP-DOWN state is considering as a dominant electrographic pattern during immobility and slow wave sleep. This study is focused on the analysis of spatial distribution and neuronal correlates of the DOWN state in rats

**Methods:** Local field potentials and multiunit discharges were recorded bilaterally in the prefrontal cortex (PFC) and hippocampus of non-anesthetized, tethered rats (Sprague Dawley, males, weight 350-400g) with 256 channel, 4 shank silicon probes. We have focused our study on the analysis of the positive wave of slow oscillations (SOs), which is considered as the DOWN (silent) state of the UP-DOWN state in the anterior cingulate (AC), prelimbic (PL) areas of PFC and hippocampus during immobility.

**Results:** Our experiments showed that SOs occurred intermittently with a mean interval 1.4±0.8 (±SD) seconds. The SOs began with the DOWN state, and they were generated locally within AC or PL areas, or simultaneously in AC, PL and hippocampus bilaterally (generalized SOs). The DOWN state of local SOs in the AC was associated with a decreased rate of multiunit discharges. Similar waves in the PL area were associated with increased multiunit discharges. We observed high speed propagation of generalized SOs that occurred with 3-6ms delay within left and right PFC and less than 10ms delay between the PFC and CA1 area of hippocampus. All generalized SOs were associated with decreased multiunit discharges.

**Conclusion:** Our data support the hypothesis that neocortical networks are sufficient to generate focal SOs but the participation of external input is needed for occurrence of generalized SOs.

## INTRODUCTION

Prefrontal cortex (PFC) is reported to participate in several different brain functions. Subregions of PFC (anterior cingulate (AC) and prelimbic (PL)) have different anatomical connections, and play different roles in brain function. Compared to PL, AC receives a significantly larger percentage of intra cortical afferents and input from anterior, as well as “relay” nuclei of thalamus ^59^, while PL receives more dense projections from the middle nuclei of thalamus ^58,59^. Functionally, AC mostly plays a critical role in memory consolidation processes ^19,20,35,41,46,47^ while PL is necessary for retrieval of fear ^4,11^.

Slow oscillations (SOs) are dominant patterns generated by PFC during immobility, slow wave sleep and light anesthesia. The two phases of SO reflect the existence of local network in a two states: the DOWN (silent) state is characterized by a suppression of the amplitude of gamma (30-90Hz) activity associated with a decrease or silence of spike generation of recorded population of neurons ^16,23,43,60^. The DOWN state is followed by the UP (active) state when the amplitude of gamma activity increases and it is associated by an increase of the rate of neuronal discharges ^8,24,44,51–53,55,60^.

Over the past 20 years the scientific community has devoted significant attention to the analysis of mechanisms of generation of the UP state, as well as its role in the process of consolidation of memory traces ^1,10,23,24,33,38,42,55^. Much less attention has been paid to analysis of propagation of the DOWN state, specifically during sleep under un-anesthetized conditions.

A goal of this study was to characterize SOs in AC and PL areas of PFC during immobility and their relation to SOs recorded in hippocampus. We carried out recordings with 4 shank silicon probes (total 256 recording sites) implanted bilaterally into the prefrontal cortex and hippocampus. We present data on the spatial distribution and neuronal correlates of the main local field potential (LFP) patterns in PFC, specifically focusing our attention on the DOWN state of the SOs.

## MATERIALS AND METHODS

These experiments were carried out in accordance with relevant NIH guidelines and regulations. All procedures were approved by the University of California Los Angeles Institutional Animal Care and Use Committee (protocol 2000-153).

### Silicon probe preparation

Silicon microprobes each containing 64 recording contacts (type 64E, consisting of 64 100 µm^2^ electrodes spaced linearly by 50 µm), were used for these experiments. The details of probes preparation were described elsewhere ^45^. Each recording site was gold plated (Sifco 80535500) with constant-potential pulses (±2.5 V relative to a Pt wire reference, 1–5 s) to reduce impedance below 0.5 Megaohms to improve the signal-to-noise ratio ^15^ using an INTAN RDH2000 electroplating board. Two 64-channel probes were glued together (Fig. 1A) and attached to both sides of a homemade micromanipulator (Fig. 1B) so that, in the anterior part of the manipulator, the distance between probes (#2 & 3) was 1mm and in the caudal part (#1 & 4) it was 8mm.

### Surgery

Experiments were performed on adult male Sprague-Dawley rats (Charles River, body weight 300-400g. Animals (n = 5) were anesthetized with 1.5–2.0 ml/min of isoflurane in 100% oxygen and fixed in a stereotactic surgery frame. Body temperature was kept between 36.6 and 38.0°C with a thermostatically controlled heating pad. The skull was opened and holes 1mm in diameter were drilled bilaterally above prefrontal cortex (AP=2.7; lateral 0.5mm from the sagittal suture), and above posterior hippocampus (AP=−2.5; lateral 4.0). After punching dura mater an array of silicon probes was attached to a stereotaxic manipulator, centered above each hole and slowly moved into the brain until the micromanipulator touched the surface of the skull. Ground and reference electrodes (stainless steel screws) were positioned in the cerebellum 2.0 mm posterior to lambda and 1.0 mm lateral from the sagittal suture. The system was fixed to the skull with dental cement. The incision was then sutured and treated with 0.25% bupivacaine as well as topical antibiotic ointment. Figure 1C illustrates a rat with implanted silicon probes connected to an amplifier

### Data acquisition

On the fourth day after surgery electrical activity was recorded in each animal’s home cage. EEG/video data were recorded with wide bandwidth from 0.1 Hz to 3.0 kHz and sampled at 10 kHz/channel using two RHD 2000 128-channel amplifier boards and data acquisition software (INTAN Technologies, LLC; Los Angeles, CA) for 6-8 hours a day, and stored on external hard drives. After completion of experiments animals were perfused transcardially with 4% paraformaldehyde (PFA) in PBS. The brain was then removed and postfixed overnight in 4% PFA in phosphate buffer at 4°C. The brains were then subjected to MRI imaging for localization of silicon probe tracks (Figure 1D).

### Data analysis

Initially the identification of the location of silicon probes was performed on ex-vivo MRI sections (Fig. 1D). The fixed brain was scanned by the Bruker 7T animal scanner (Brain Research Institute, UCLA), using a 14mm diameter 1H radiofrequency coil. The 3D T2 RARE sequence with parameters of TR/TE = 1700/63 ms, 200×128×128 with voxel size 80 µm^3^ and RARE factor = 6 was applied. Gamma event coupling analysis, described in our previous publications ^3,26^, was performed for more precise location of recording sites within PFC and hippocampus (Fig. 1E). It identifies the local networks and allows distinguishing the location of recording sites in the AC, and PL areas of PFC as well within different areas of hippocampus. In addition, current source density analysis (CSD) of the sharp wave-ripple complex identified layers within the CA1 area and dentate gyrus (DG) ^7,61^. Specifically, the discrete second derivative across the depth electrode sites was computed on the averaged local field potentials (LFPs) with two-site spacing to reduce noise (Bragin et al., 1995, 1997; Ylinen et al., 1995). The data were then further smoothed with spatial interpolation to produce the CSD map. Current sinks and sources associated with sharp waves and ripple oscillations in normal conditions provided precise landmarks for the identification of the recording sites located within pyramidal layer of the CA1 area and was extrapolated to identify layers in DG. In addition, unitary activity in the CA1 pyramidal layer provided further help for the depth calibration of the electrodes within hippocampus.

**Figure. 1.**
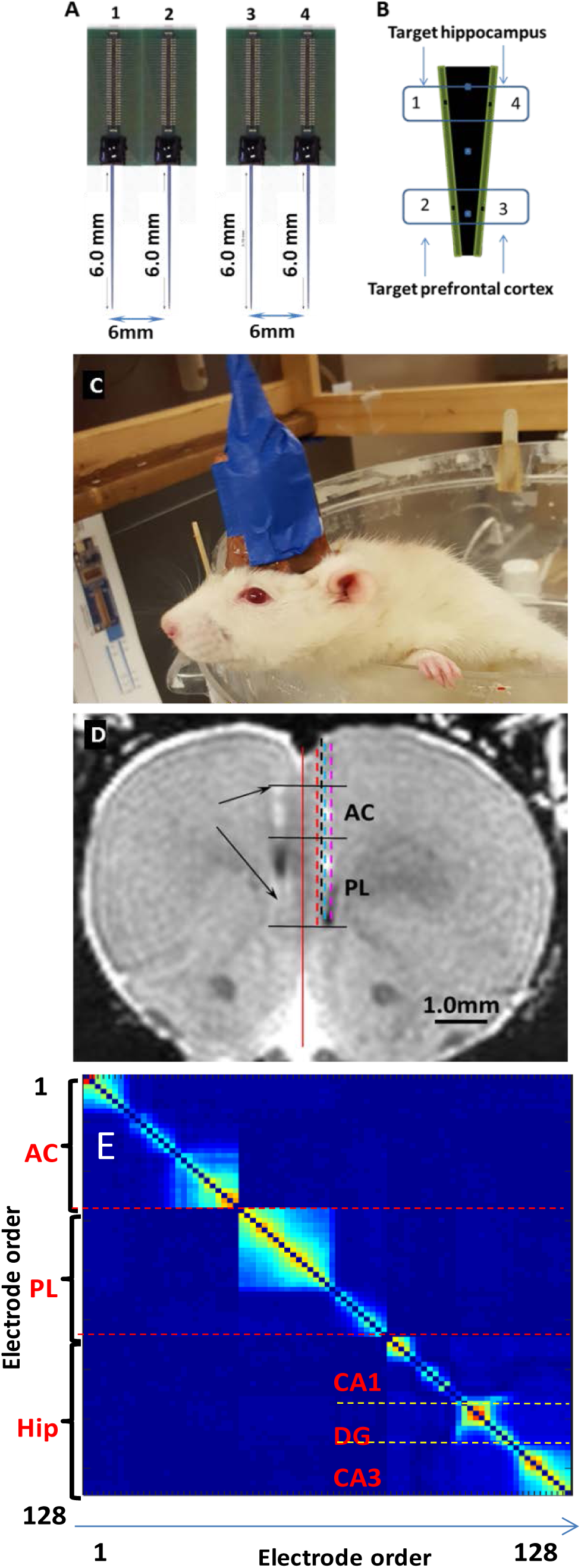
A schematic of the silicon probe arrangement for implantation. A – two pairs of glued silicon probes. Numbers indicate the length of probes and distance between shanks. B –arrangement of silicon probes on the micromanipulator and target areas for each of them. C – a picture of a rat with implanted the silicon probe system. D – A section of an MRI image with the track of the silicon probe on the left (arrows) and right sides from one of the rats. Dashed lines on the right side are schematic representations of silicon probe tracks from other four rats. E – identification of the location of recording sites by gamma event coupling. Abbreviations: AC and PL correspondingly indicate anterior cingulate and prelimbic areas of PFC. Hip – hippocampus; CA1, CA3 and DG – correspondingly CA1, CA3 areas of hippocampus and dentate gyrus.

Data were then reviewed on the computer screen, and one hour files were selected from each animal, when it was immobile. The quality of selected files was verified by power spectrogram analysis of electrical activity recorded from the CA1 areas of hippocampus. If the spectrogram contained the peak at the frequency band 3-10Hz, which characteristic of exploration activity or rapid eye movement sleep ^5,6^, they were rejected from the analysis. During this immobile periods animals may be sleeping, and because we did not performed a sleep scoring, we will refer the selected files as recorded during the immobility.

Datapac 2K2 software (RUN Technologies, LLC) was used for detection of SOs. Initially four recording sites located in the middle left and right AC and PL areas were selected. After low pass 2Hz filtering (FIR, order 221) SOs exceeding 3 standard deviations (SD) were detected and then reviewed for artifacts. SOs that occurred in all 4 recording sites were considered as generalized SOs, and those that occurred in any single recording site were considered as local SOs. They formed different buffers for further analysis. For analysis of the delay of occurrence of SOs in different brain areas, generalized SOs were normalized and time difference between selected SOs was measured in the middle of the ascending slope of SOs using Datapac 2K2 software. We calculated a modified shift predictor for the event synchrony between left and right AC and PL, emulating procedures described in previous studies ^36,49^. First, the distribution of delays from all measured events was calculated. Second, the random events were generated by assigning an equal number of events with bootstrap sampling of 1000 times. Third, a normalized synchronization metric was computed by summing the measured values in a 1-3 ms window and then dividing by the corresponding area of the chance distribution. We then calculated whether the incidence of delays less than 3ms was greater or equal to the chance.

Multiunit discharges (MUD) of neuronal activity were selected after high pass 600Hz filtering and setting the threshold at 2 SD. For voltage versus depth analysis all detected events were averaged and values of the peak amplitudes were measured at each recording site. The values were normalized and plotted against each recording site.

#### Statistics

General descriptive statistics were performed to summarize the mean and standard deviations of two types of SOs. Histograms were created to describe the distributions of inter SOs intervals in the datasets. All the statistical analyses that performed in the current study were based on a significant criterion of p<0.05.

## RESULTS

Analysis of silicon probe tracks revealed that recording sites were located within layers 3-5 a distance of 2mm within AC and 1.5 mm within PL (see Figure 1D as example).

During the slow wave period, SOs occurred intermittently (Figure 2A) at 0.3 - 5.0 second intervals (Mean 1.4±0.8/sec,[±SD]), (Figure 2B). Two types of prefrontal cortex SO were observed in our experiments: (1) local with maximum amplitude either in the AC or in the PL; (2) generalized, which occurred simultaneously and bilaterally in the AC and PL areas of the prefrontal cortex, and in the hippocampus.

**Figure 2.**
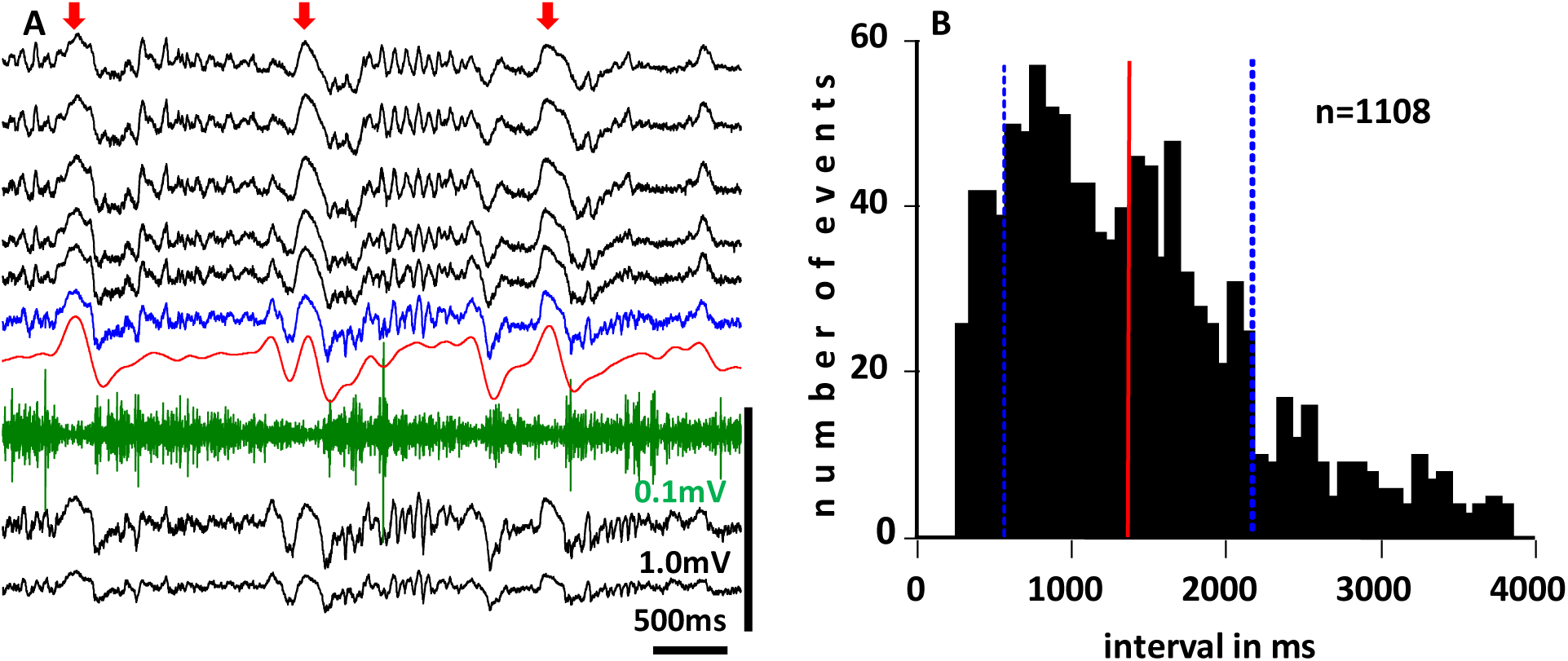
A – an example of SOs (red arrows) recorded in the AC, low pass filtered data (red line), and multiunit activity (green line) recorded during slow wave state. B – Distribution of inter SO intervals recorded during selected 75 minutes of slow wave periods from 5 rats. A total 1108 SO events were quantified. The red line indicates the mean and blue lines +/− standard deviation (SD) of inter event intervals at 791.75 and 1401.23 ms.

### Local slow oscillations

Two types of local SOs were observed in the PFC. The first type, which constituted 24% of all SOs, had maximum amplitude in the AC area of prefrontal cortex. The second type, which constituted 11% of all SOs, had a maximum amplitude in PL (Figure 3,A,D). Figure 3C illustrates voltage –depth profiles of SOs originated in the AC (black lines) and in the PL (blue lines). Both types of SOs occur intermittently either within AC or PL areas without interference with each other. Thirty percent of AC’s SOs were followed by spindle like activity, while the PL SOs were never associated with spindle like activity. The rate of MUD, recorded in AC decreased during the AC’s SOs. In PL at the same time a partial suppression of MUD was observed in the upper part, and there was no significant change in MUD recorded in multiple locations in the lower part of PL (Figure 3B). During PL’s SOs there were no changes in rate of MUD in multiple areas of AC (Figure 3, E, top part). Prior and during the ascending phase of SOs there was an increase in MUD rate in the upper part of the PL, while there was no significant change in the lower part (Figure 3, E).

**Figure. 3.**
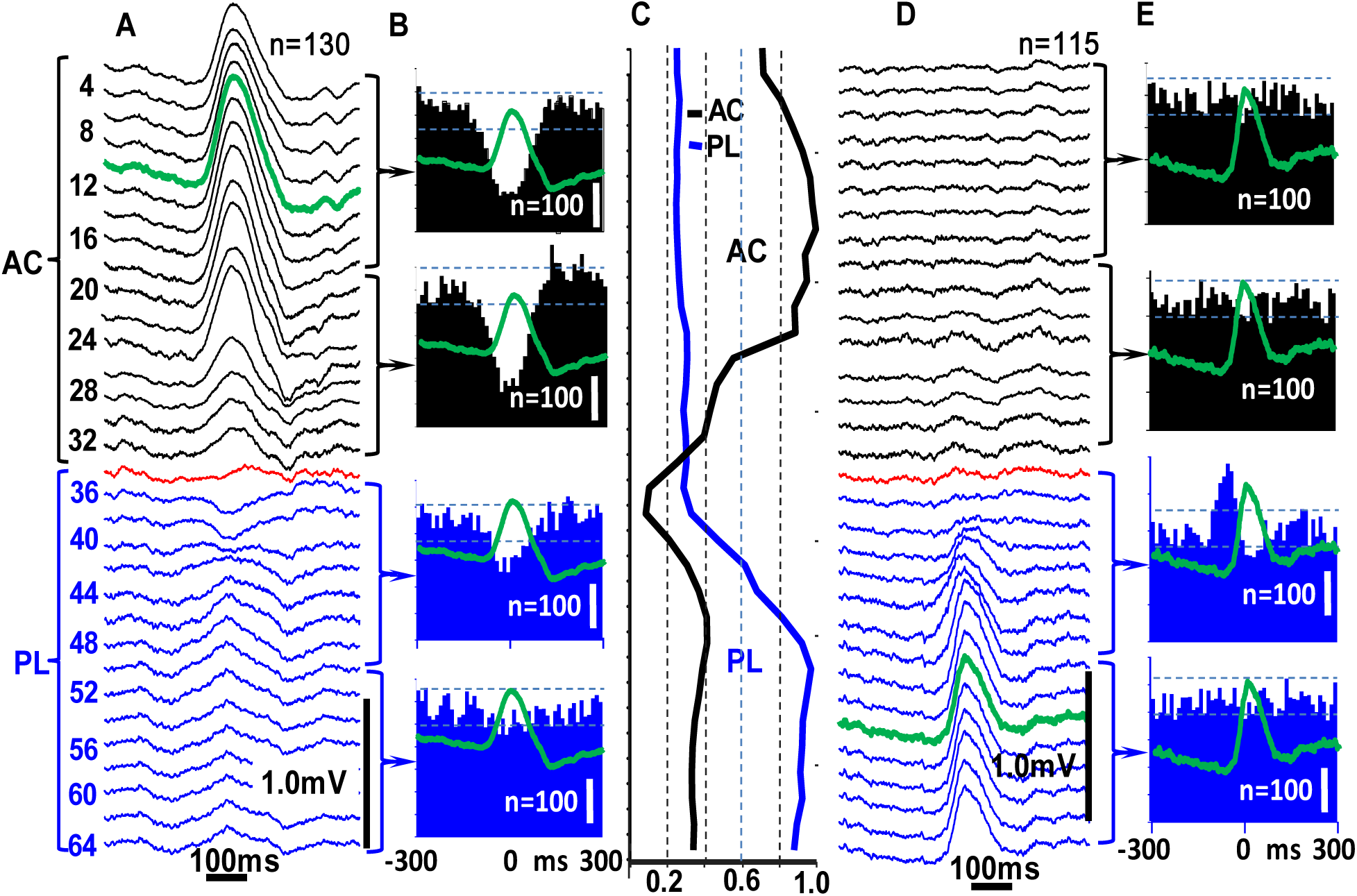
Spatial characteristics of SOs in the prefrontal cortex. (A) Voltage-depth profiles of SOs (n=130) with maximum amplitude in the AC and (D) SOs (n=115) with maximum amplitude in PL. Red line indicates the border between anterior AC and PL cortices. B - synchrony of MUD during AC SOs and E - synchrony of MUD during PL SOs. Each histogram represents summated MUD from four recorded sites within the area indicated by the bracket. C – graphic presentation of changes of amplitude versus depth profiles of the AC SOs (black) and the PL SOs (blue). Y-axis – number of recording sites, X-axis - normalized amplitude of events. Dashed lines indicate 95% of confidence interval. “n” inside histograms indicates calibration for 100 spikes and on the top the number of averages.

### Generalized SOs

Generalized SOs were observed in 65% of all recorded SOs. They occurred simultaneously in left and right PFC as well as in the hippocampus (Fig. 4) and consist of positive waves with a mean duration of 280±70ms (±SD). In the CA1 region of hippocampus the SOs consisted of only by a positive wave only in the CA1. In the hilus of the DG a negative wave preceded the CA1 positive SOs (Figure 4, C and Figure 5 B). Current source density analysis indicated that the main source for this early wave was located in the stratum moleculare of the CA1 and DG (Figure 5C, arrow), which is a source of projections from entorhinal cortex.

**Figure 4.**
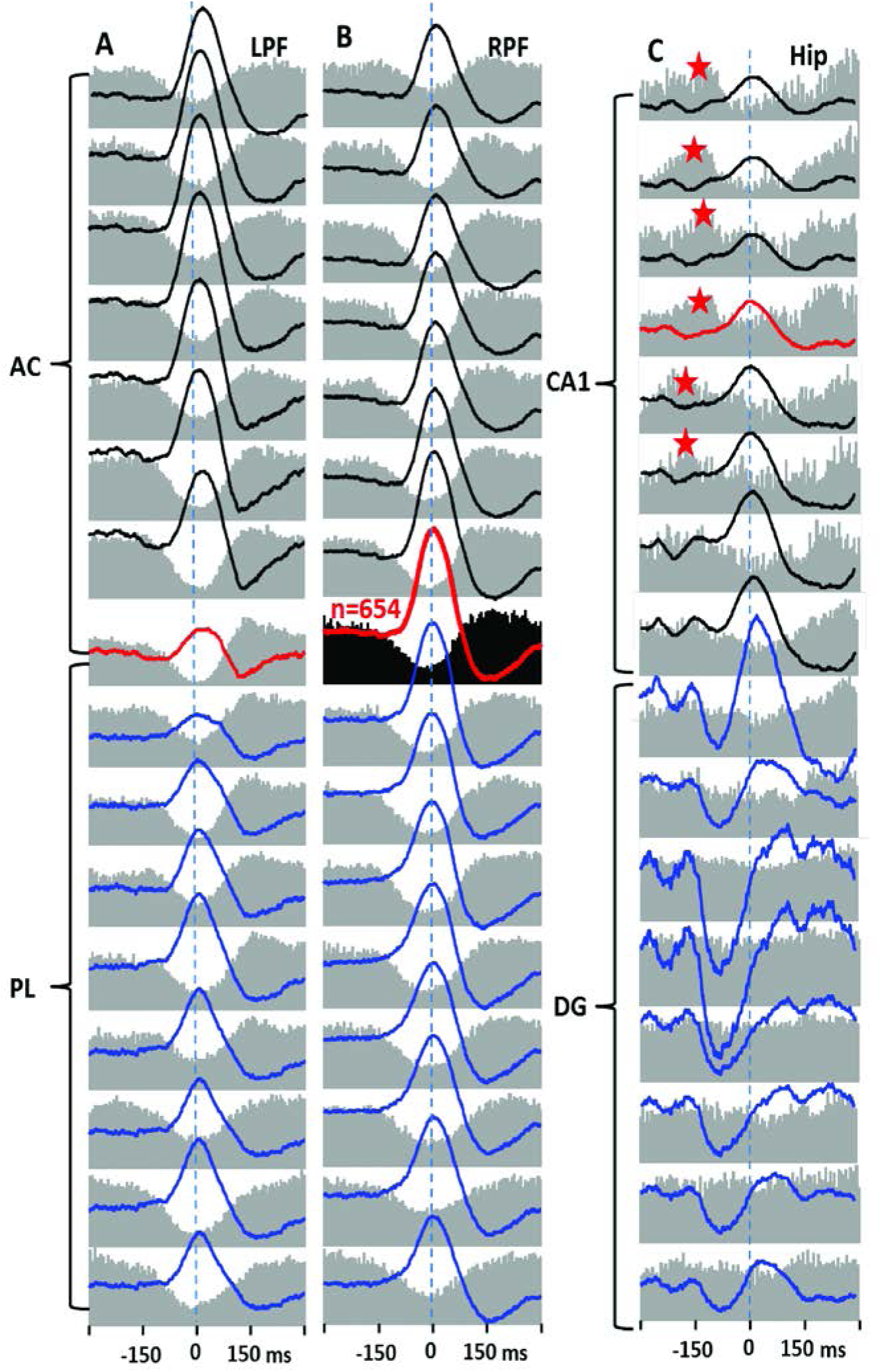
Generalized SO in the left and right PFC and in the right hippocampus. Examples of averaged SO (n=654) recorded with the distance of 100µm are presented in each column. For the averaging trigger, the SO recorded at the border between the right anterior cingulate (AC) and prelimbic (PL) areas was chosen. Perievent histograms of multiunit discharges (MUD) are presented in grey behind every SO. The red lines in A and B indicate the border between AC and PL and in C - the pyramidal layer of the CA1 area of hippocampus. Dentate gyrus (DG) is labeled by blue color. Horizontal dashed lines indicate 95% confidence intervals

**Figure 5.**
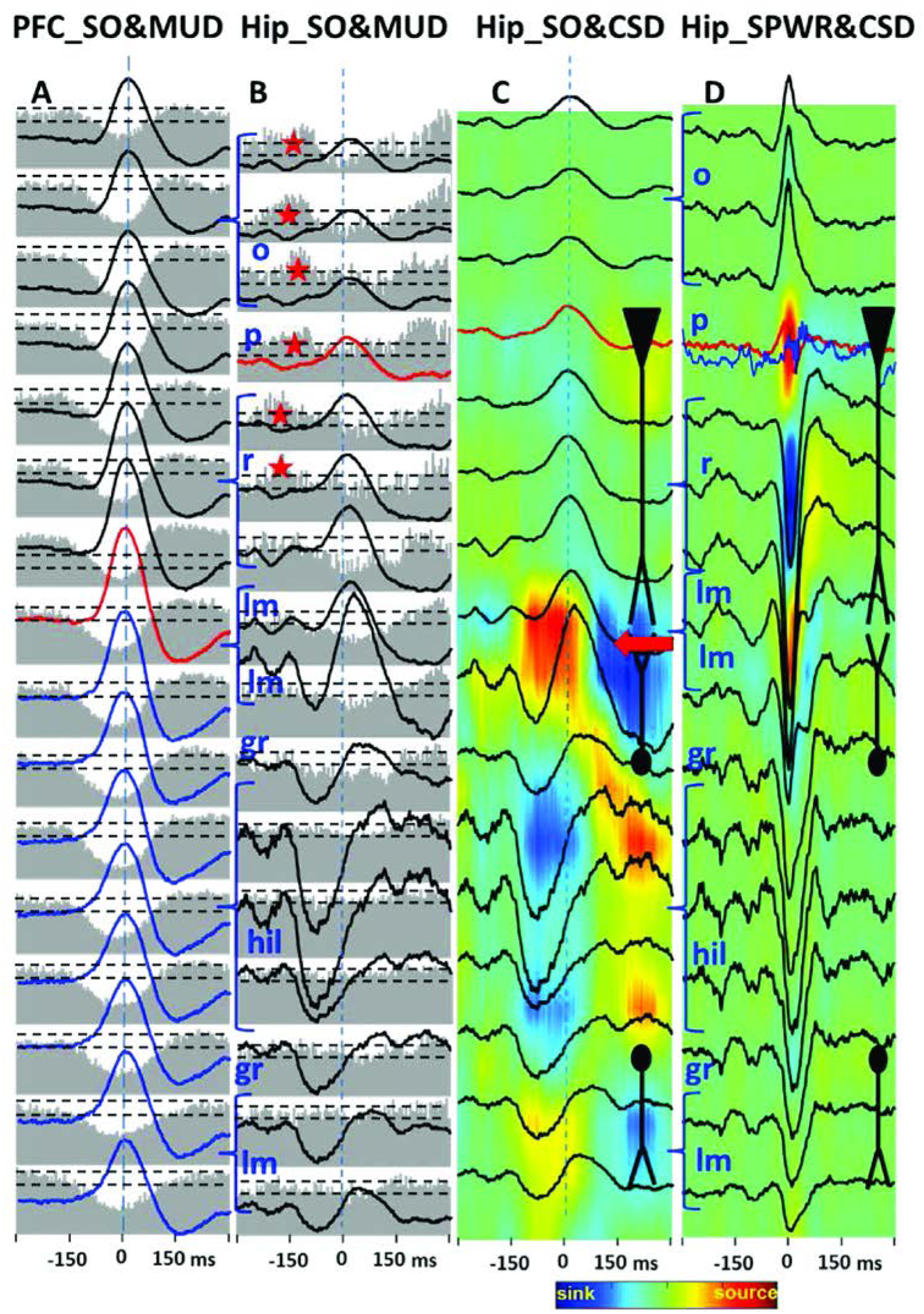
Current source density analysis of SO recorded in the hippocampus. A and B are copies of averaged SOs and MUD histograms from the figure 3. C – current source density analysis of hippocampal SO. D – current source density analysis of sharp wave – ripple (SPWR) recorded in the same file. Abbreviations: o, p, r, lm, gr, hil – correspondingly stratums orience, pyramidale, radiatum, lacunose-moleculare, granular layer of DG and hilus. See details in the text. Red arrow indicates the source of negative SO in the DG. Horizontal dashed lines on A and B indicate 95% confidence intervals.

In both areas of prefrontal cortex these generalized SOs were associated with suppression of multiunit discharges (MUD) (histograms in the Figure 4,A & B, 5A). This suppression of MUD was more prominent close to the border between AC and PL areas. In the hippocampus the suppression of MUD was less prominent, was observed only in the CA1 (Fig. 4C, 5B) and was preceded by an increase of MUD associated with the negative wave in the hilus of the DG (Fig. 4C, 5B red stars). There were no significant changes in MUD in the DG during the DOWN state (Figure 4C, 5B), where this preceded the negative wave, which began in average 96±12ms earlier. However average events may be misleading because they may just show variation of mean values as a result of prominent variation of individual values (Figure 7a). More detailed analysis of synchrony of the DOWN state occurrence was performed on individual records from left and right prefrontal cortex, recorded between left and right areas of PFC as well in the CA1 showed that in all brain areas SOs occurred with a delay no longer than 10 ms (Figure 6, superimposed traces and the inset). Sixty seven percent of all generalized SOs occurred within a 10 ms time window.

**Figure 6.**
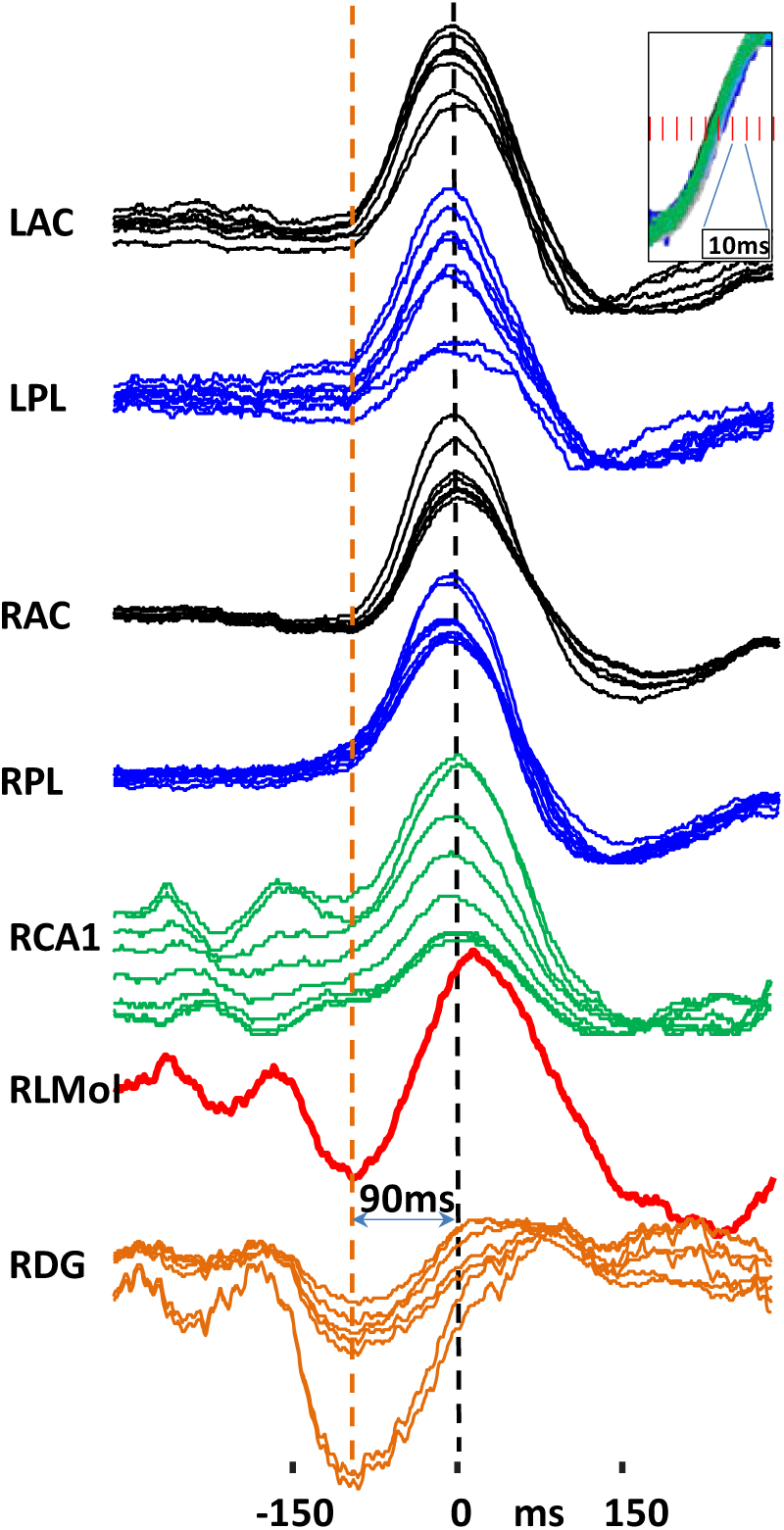
Superimposed generalized SOs (n=84) in the left and right PFC and in the right hippocampus from every 4^th^ recording site. Dashed lines indicate the maximum amplitude of SO recorded in the DG (orange) and maximum amplitude of SO recorded in the left and right prefrontal cortex and hippocampus (black). The inset illustrates the superimposition of normalized slops in all areas of PFC and in the CA area of hippocampus. RLMol – right lacunose moleculare area in CA1.

**Figure 7.**
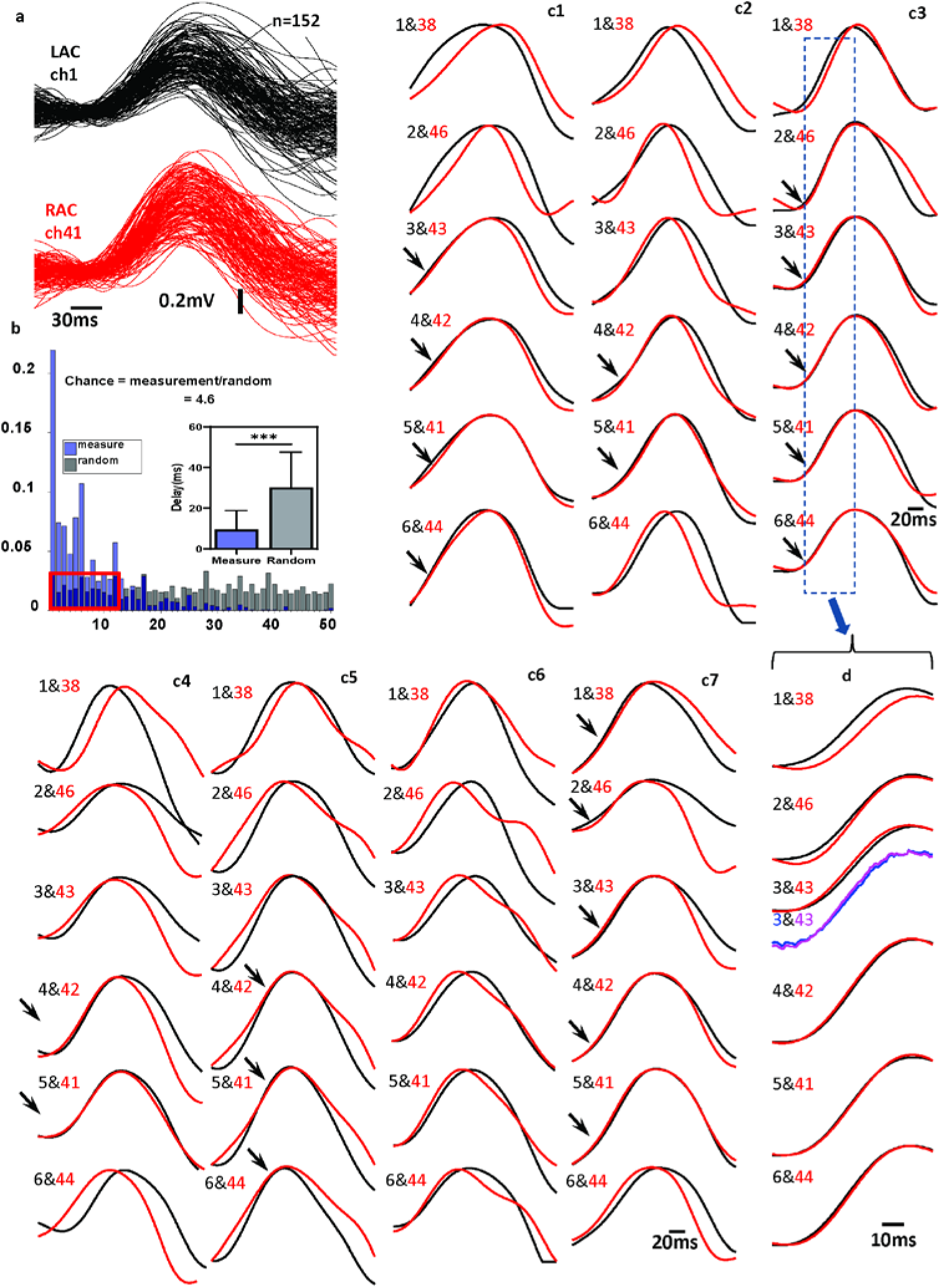
Variation of individual SOs and their synchrony between left and right anterior cingulate cortex. **a**. Superposition of 152 SOs recorded in the left (black lines) and right (red lines) anterior cingulate cortex. **b** – Blue bars distribution of delay of generalized SOs between eight recording sites located in the left and right AC, black bars distribution after bootstrapping (red box indicates the level of coincidence by chance). The inset is the graphic representation of the measured delays, which is significantly shorter (p<0.001) in comparison to randomly selected events. c1-c8 examples of individual SOs recorded in these areas with recording sites separated by 200µm. Arrows indicate those events where time difference between left and right sites were less than 10ms. **d.** Ascending slope of SO b4. Recoding sites 3&43 colored as blue and purple are the same events as red and black, but recorded relatively to the reference electrode located in the thalamus.

However average events may be misleading because they may just show variation of mean values as a result of prominent variation of individual values. More detailed analysis of synchrony of the DOWN state occurrence was performed on individual records from left and right prefrontal cortex, where we calculated delay of individual DOWN events. Examples of these events are presented in Figure 7a, were variability of slopes of ascending and descending phases is more than 30 ms. However, in 20-40% of individual pairs (mean 29.7%) recorded in the left and right sides the slopes of the DOWN state occurred with a delay of 1-3 ms (Figure 7 c1-c7, arrows). This almost simultaneous occurrence does not depend on the reference electrode, because it occurs only between some recording sites, but not between others, and it has a similar delay even in cases when it was recorded relative to other reference electrode (Figure 7 d, recordings 3&43 red and black were recorded relative to one reference electrode and purple and blue relative to another reference electrode). We calculated a modified shift predictor for the event synchrony between left and right AC and PL. The slope delays from a total of 1158 events were measured, and the random events were generated by assigning an equal number of events with bootstrap sampling of 1000 times. The histograms in Figure 7b show the normalized probability of the measured event delays (blue bars) and the bootstrap random sampled data (black bars). The occurrence of SOs within a 1-3 ms window is 4.6 times higher than chance.

To evaluate the synchrony of occurrence of SOs, we further estimate the SOs peak latencies between left and right AC areas in six different paired recording sites (named ch1 – ch6). The one-way ANOVA for the latency data indicated no statistically significant differences between LAC and RAC in all 6 selected recording sites (F (5, 876) = 1.38, p = 0.23, Figure 8). Detailed results are summarized in the table 1. Our data indicated that the latency of SOs between LAC and RAC shows no changes in response to different recording areas, where we calculated delay of individual DOWN events. Examples of these events are presented in Figure 7a, were variability of slopes of ascending and descending phases is more than 30 ms. The histograms in Figure 7b show the normalized probability of the measured event delays (blue bars) and the bootstrap random sampled data (black bars). The occurrence of SOs within a 1-3 ms window is 4.6 times higher than chance.

**Table 1.** Time delays between SOs recorded in 6 recording sites of the left (LAC) and right anterior (RAC) cingulate cortex.

|  |  | LAC-RAC<br>latency<br>(Mean) | LAC-RAC<br>latency<br>std) | Number<br>of events |
| --- | --- | --- | --- | --- |
| Recording Sites | Ch1 | 8.86 | 8.42 | 147 |
|  | Ch2 | 7.78 | 8.44 | 147 |
|  | Ch3 | 8.83 | 7.50 | 147 |
|  | Ch4 | 7.66 | 7.26 | 147 |
|  | Ch5 | 7.81 | 7.50 | 147 |
|  | Ch6 | 9.60 | 9.40 | 147 |

**Figure 8.**
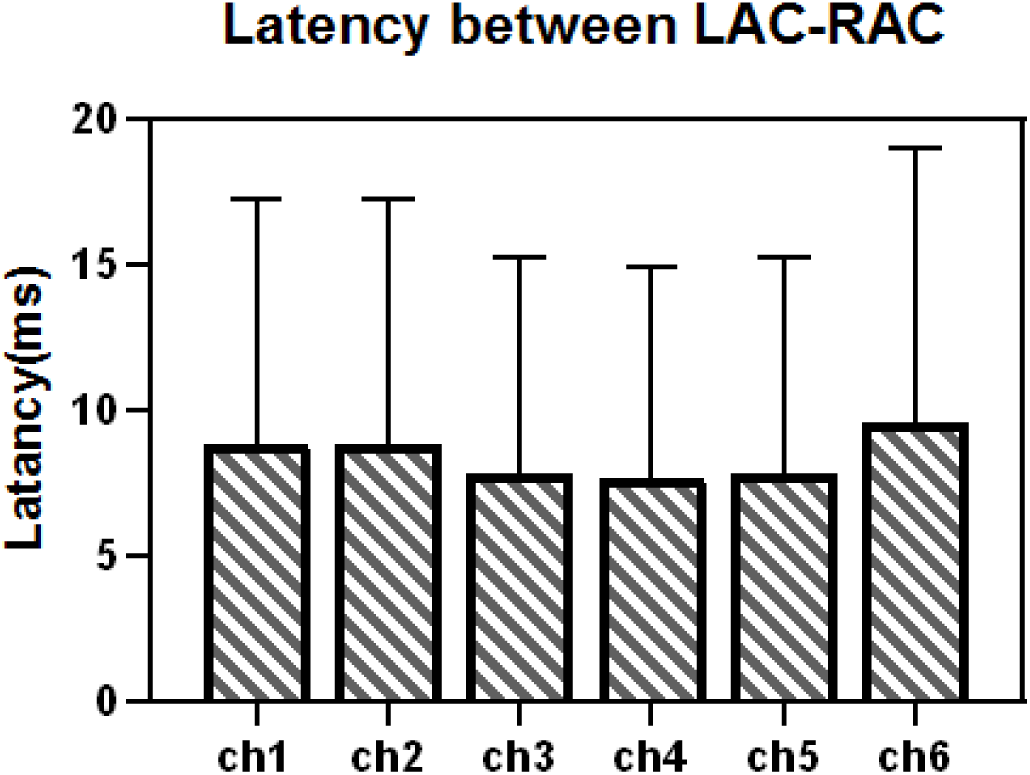
Comparison of the SOs peak latency between LAC and RAC in a total 882 pairs from six different recording sites. No statistically significant differences

## DISCUSSION

The main finding of this study is a demonstration that during immobility generalized SOs occur intermittently and begin with the DOWN state associated with the suppression of MUD. This DOWN state occurs within 1-3ms delay bilaterally within subregions of PFC and hippocampus. Local similar SOs recorded in the PL area are associated with an increase of MUD.

Several publications showed that SOs is a wide spread electrographic pattern involved both neocortical and hippocampal areas of the brain ^24,30,40,52,55^. Our findings extend these data by showing that this wide spread pattern begins with an initial DOWN state. It does not contradict the existing hypothesis of wide spread excitatory front of UP state ^24^, and suggest that this initial UP state occurs as a rebound after an initial DOWN state

According to numerous publications [see reviews ^12,13,33^], during the awake state and rapid eye movement sleep the membrane of neurons is slightly depolarized. The electrophysiologically visible UP-DOWN state in the neocortex occurred when depolarizing influences from the sensory systems and brainstem areas decreased.

It is hypothesized that the DOWN state has occurred as a result of disfacilitation and temporal absence of synaptic activity ^2,9,50,56^. Usually the occurrence of DOWN state is considered as a process of recovery after termination of the UP state ^27,43,55^. The possible mechanism of the DOWN state as recovery process after the UP state is turning on of the hyperpolarization activated low threshold CA^2+^ and Ih currents ^28,32,48^ and occurrence of miniature EPSPs ^55^. This may be the case in recordings during deep slow wave sleep, under anesthesia and in vitro experiments. In our experiments we have investigated pattern and neuronal correlates of SO during immobility and light sleep where SOs occurred intermittently with the rate 1.4/sec. These intermittently occurring SOs began with the DOWN state indicating that the DOWN state may occur initially form the baseline activity and is followed by the UP state. We cannot exclude the possibility that these initially occurring DOWN state has the same mechanisms it was proposed in previous studies, however it is not a recovery from the UP state. The absence of the increase the synchrony of MUA before the DOWN state allow to suggest that DOWN state may occur as a result of initial inhibition of the neuronal activity. ^16^ showed that specific type of interneurons (GIN cells) fired persistently during the DOWN state. So the DOWN state could be an initial step in the occurrence of repetitive of UP-DOWN state activity.

### Local SOs

Demonstration of local SOs in different areas of neocortex is in accordance with many previous data indicating the existence of local SOs and support the hypothesis that local neocortical networks are sufficient to generate SOs ^23,30,40,43,54,60^. In the AC local DOWN state involved neuronal populations along 2mm of this area in the layers 3-5 and was associated with suppression of MUD during the ascending and descending phases of SOs. At present, mechanisms triggering the DOWN state remain unclear. ^27^ described an occurrence of long (100-300ms) IPSPs in the period about 250 ms before the onset of the DOWN state, which suggests the existence of interneurons, that trigger the silent period within a surrounding network. Potentially, it may occur with involvement of GABAa and GABAb receptors ^29^ or D(1) -like dopamine receptors ^31^. Recently two groups showed that inhibitory somatostatin containing interneurons (SOM-Ins) in the superficial layers of neocortex, which may be are Martinotti neurons ^17^, play a key role in moving the network into the silent DOWN state. ^21^ found that chemogenetic activation of SOM-Ins increases slow wave activity. ^39^, showed that during the transition into the SOs DOWN state SOM-Ins, but not parvalbumin-positive interneurons or pyramidal cells, increase frequency of discharges. Martinotti cells may form an electrically coupled network that exerts a coherent inhibitory influence on its targets and to play a role as "first responders" when cortical excitatory activity increases ^17,25^. However, similar local positive waves in the PL area are associated with an increase in MUD. Mechanisms of generation of this type of wave are also unclear.

### Generalized SOs

Studies from Steriade’s Lab showed that SOs in the cat neocortex could be synchronized along on the distance of 7 mm ^14,57,60^. Generalized SOs are considered as propagated waves from frontal to occipital parts of the brain ^27,30,44^. The speed of propagation varies in different publications. It is 1.2–7.0 m/sec in humans ^30^ and a much slower 0.4m/sec in rodents ^44^. In some publications ^30,34^ simultaneous occurrence of the DOWN state was mentioned, but no quantitative analysis was performed. The corpus callosum may be the pathway that synchronizes SOs, because Mohajerani et al., ^34^, using the technique of voltage sensitive dye imaging, showed that synchrony of SOs decreases in acallosal mice.

In our experiments, in all investigated brain areas: left-right AC, PL and CA1 area of hippocampus the DOWN state of generalized SOs occurs within a 10ms window, and 1-3 ms between left and right AC and PL. If we take into account the maximum speed of propagation for the UP (active) state indicated by Massimini et al.,2004^30^ in human (7.0 m/sec) it may suggest that the DOWN (silent) state of SOs generated in the PFC could propagate to hippocampus within the 10 ms window. However, if the speed of the DOWN state of SO propagation in rats is similar to that for the UP state in mice ^44^, than we should see a delay between the occurrence of SOs in the PFC and hippocampus. Moreover, the DOWN states recorded in the left and right hemispheres, in a significant number of cases, much higher than chance, occurred with a 1-3msec delay, which is shorter than latency of propagation of electrical activity via corpus callosum. The synchrony of UP-DOWN states observed under anesthesia or in in-vitro slice preparation experiments, where SOs occurred rhythmically the synchronization of DOWN state, might be explained by a synchronous release of DOWN state from the UP state. However, as we described in our experiments, during immobility SOs occur intermittently and begin with the DOWN state. It cannot be a simultaneous release from the UP state and the initial inhibitory process should be involved in generation of intermittent SOs. These data indicate the existence of a common trigger, most likely in subcortical areas, which synchronize the occurrence of generalized SOs. This hypothesis was suggested by ^60^ and was confirmed in ^22^ experiments. This common trigger, may interfere with local mechanisms triggering the DOWN phase and synchronizing their simultaneous occurrence of SOs in multiple brain areas. It was shown that D1-like receptors are essential for the occurrence of SOs ^31^, so a potential candidate for this role could be substantia nigra, which has diffuse projections to multiple brain areas. Another potential candidate could be the central lateral thalamic nucleus, which is as a key region in the subcortical arousal systems for maintaining the level of consciousness ^18,37^. Both of these areas could synchronize local neocortical networks.

In conclusion, the results of our study have raised a number of questions for future experiments. Among them are unravel the mechanisms for simultaneous entry into the DOWN state in multiple brain areas, and what is the functional role of local SOs in the AC and PL cortices.

## Acknowledgements

This research was funded by NIH NINDS grants NS065877 (AB), NS33310 (JE), and NSF Neuronex grant 1707408 (SCM).

## Author Contribution statement

Dr. A. Bragin, J. Engel Jr. and S. Masmanidis desinged experiments. Dr. S. Masmanidis provided silicon probes, Dr. L. Li and Dr. Bragin performed experiments and data analysis. All co-authors participated in discussion of the results and writing of the manuscript.

